# The extended second law of thermodynamics suggests a thermodynamic imperative driving the evolution of life systems towards increased complexity

**DOI:** 10.1101/2021.09.13.459895

**Authors:** Sergio Menendez

## Abstract

From a thermodynamic point of view life structures can be viewed as dissipative systems capable of self replication. Energy flowing from the external environment into the system allows growth of its self replicative components increasing the system complexity concomitantly with an increase in the entropy of the universe, thus observing the second law of thermodynamics. However, general thermodynamic models of life systems have been hampered by the lack of precise equations modelling far from equilibrium driven systems operating in non-linear response regimes. Recent theoretical advances, applying time reversal symmetry and coarse grained state transitions, have provided theoretical insights into the thermodynamic constraints that bind the behaviour of such far from equilibrium life systems. Setting additional constraints based on empirical observations allows us to apply this theoretical framework to gain a further semiquantitative insight on the thermodynamic boundaries and evolution of complex self replicative life systems. This interpretation suggests a thermodynamic hierarchical organisation based on increasing accessible levels of usable energy, which in turn drives an exponential punctuated growth of the system’s complexity. For the earth life system this growth has historically not been limited by the total energy available from the external driving field, but by the system’s internal adaptability needed to access higher levels of usable energy. Therefore, in the absence of external perturbations, the emergence of an initial self replicative dissipative structure capable of variation that enables access to higher energy levels is sufficient to drive the system’s growth irreversibly towards increased complexity across time and space in a hierarchical manner. This interpretation is consistent with current empirical observation of life systems across both time and space and explains from a thermodynamic point of view the evolutionary patterns of complex life systems on earth.

## Introduction

A consensus definition of what features define a life system remains controversial (Schrödinger, 1967). However, among the basic requirements for life the capability of a system to self replicate and therefore propagate itself over space and time seems to be accepted as a minimal essential feature. Self-replicative (SR) live systems appeared on earth around 4.2 billion years ago. Current hypotheses speculate that polymeric assemblies (such as RNA) emerged under suitable environments as the original SR systems, however the precise mechanisms and environment required for this event remain elusive (Attwater et al., 2010; Nakashima et al., 2018; Walker, 2017). From a biochemical point of view the process of abiogenesis could be viewed simplistically as a reaction/s where a number of basic components (reactants) organise themselves into more complex assemblies (product/s) that acquire the emergent property of self-replication (SRn). Whether an external energy source was needed to catalyse this initial step, or thermal fluctuations within the equilibrium frame of an existing reversible reaction were sufficient to acquire a higher energy level assembly capable of SRn remains unclear (England, 2015). Nevertheless, in order for there to be further growth, the original SR assemblies would have required to be able to tap into an external energy source to drive future self-replication and escape further from equilibrium (Duim & Otto, 2017; Prigogine & Nicolis, 1971; Ragazzon & Prins, 2018). The nature of this original accessible energy source and the initial equilibrium conditions remain unknown, though several hypotheses have been proposed (Walker, 2017). Thus, from this thermodynamic perspective, live structures subjected to an arbitrarily time varying external driving field (external energy source) can be viewed as dissipative open systems capable of self-replication (Prigogine & Nicolis, 1971; Ragazzon & Prins, 2018). Several basic “close to or at equilibrium” models of live structures have been proposed (Kleidon et al., 2010; Prigogine & Nicolis, 1971; Spanner, 1953). However, the statistical behaviour of far from equilibrium life systems cannot be modelled based on the properties of their individual microstates at one moment in time alone and, thus, these models are limited in the real world. Furthermore, efforts to derive a more general thermodynamic theory of life systems have been hampered by the lack of precise equations for far from equilibrium systems subjected to arbitrarily time varying external driving fields, as these systems operate in a non-linear response regime that is difficult to model using classical statistical thermodynamics (Crooks, 1999; England, 2013). Recent theoretical advances focusing on comparing dynamical trajectories, rather than describing local microstates, have made further progress. By applying time reversal symmetry and coarse grained state transitions concepts, these studies have provided helpful insights into the thermodynamic constraints that bind the behaviour of such self replicative systems (Bartolotta et al., 2016; Crooks, 1999; England, 2013; Jarzynski, 2011; Pinero & Sole, 2018). The great advantage of these models is that they are derived under general premises and hold even when an arbitrarily time varying external field drives the system in far from equilibrium conditions. Thus, this theoretical framework, generally termed Extended/Bayesian Second Law (ESL) of thermodynamics, allows us to study systems subjected to an external field driving undergoing an irreversible reaction of interconversion between two arbitrarily chosen coarse grained states defined by some feature of the system that can be monitored. Importantly, complex life biological systems can be studied in this manner and the theoretical background validating this application has already been demonstrated (Bartolotta et al., 2016; Crooks, 1999; England, 2013 and 2015; Jarzynski, 2011; Pinero & Sole, 2018).

Interestingly, using these models it is possible to evaluate minimum boundary thermodynamic values that constrain a set of arbitrarily selected initial and final coarse grained states in relation to their trajectory probabilities (equation 1). Or in other words, evaluate how the reversibility of an interconversion reaction between two arbitrarily selected coarse grained states is limited by thermodynamic boundaries determined by the amount of work available to the system and the changes in internal energy and internal entropy between the states, and vice versa. In the original 2013 England paper detailing this approach it was shown how values for the terms in the ESL equation could be calculated for irreversible SRn reactions of simple biological system such as RNA, DNA and single bacterium replication (England, 2013), demonstrating, by applying this model to irreversible reactions where the complexity of the system increases in the form of both internal energy (increase) and internal entropy (decrease), that changes in complexity must be accompanied by work made available to the system of equal or higher value to a minimal boundary value set by the irreversibility of the reaction (probability of the forward over the reverse trajectory). Thus, for inherently irreversible reactions such as the SRn of a life biological system, a minimal boundary value of available external work directly proportional to the reaction irreversibility is needed to drive changes in the complexity of the system (England, 2013 and 2015).

This observation had intuitively already been appreciated in the study of more complex ecological systems and several efforts to develop a conceptual framework for its thermodynamic analysis have been historically put forward. This led to the development of concepts such as Minimum Entropy Production, Maximum Entropy Production, Maximum Exergy Production, etc (Jorgensen, 2005; Jorgensen et al., 2007; Kleidon et al., 2010; Meysman & Bruers, 2010; Silow & Mokry, 2010; Spanner, 1953). Furthermore, field studies in complex ecological systems designed to calculate experimental values for thermodynamic-based ecological goal functions have been carried out (Fath et al., 2001; Holdaway et al., 2010; Jorgensen, 2007; Meysman & Bruers, 2010), yielding results in agreement with this intuitive understanding of the thermodynamic constraints underpinning life systems. Nonetheless, up until the publication of a formal ESL model (Bartolotta et al., 2016; England, 2013; Jarzynski, 2011; Pinero & Sole, 2018), a satisfactory theoretical framework to incorporate these concepts into a coherent ecological model supported by experimental observations had been lacking, mainly because complex ecological systems operate at the far from equilibrium conditions referred to above and using classical thermodynamics analysis of microstates could not be used to model these conditions. Importantly though, we have now at our disposal a thermodynamic theoretical framework that can be used to study the evolution of complex life systems across time and space and on this work we propose a small set of empirically self evident axioms that allow us to further expand our insight into the thermodynamic behaviour of such systems using the Extended/Bayesian Second Law of thermodynamics.

### ESL based semiquantitative analysis

As explained above, the Extended/Bayesian Second Law (ESL) sets minimum reaction thermodynamic boundary values for the conversion between arbitrarily chosen initial and final coarse grained states 1 and 2 (CGS1 to CGS2) based on the probabilities of the forward and time reversed symmetry reactions:

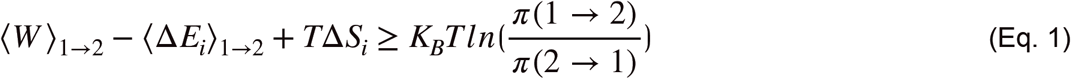

Or (in this manuscript concerning self replicators: SR)

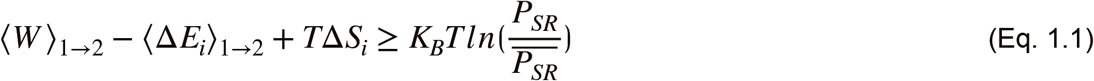

Where 〈*W*〉_1→2_ represent the average work entering the system from the external driving field, 〈Δ*E*〉)_1→2_ the average variation in internal energy of the system, *T*Δ*S_i_* the change in internal entropy of the system and 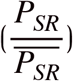 the probability of the forward self-replication reaction divided by the probability of the time reversed symmetry reaction. *T* stand for temperature and *K_B_* for the Boltzmann constant.

This equation (eq.) was derived under very general conditions and can be applied to a range of settings, including far from equilibrium driven systems (Bartolotta et al., 2016; England, 2013, 2015; Pinero & Sole, 2018). Importantly, the initial and final coarse grained states that define the terms of the equation can be selected arbitrarily to fit any desired conditions. Furthermore, as previously justified (England, 2013), when applying this equation to the process of self-replication, we can intuitively assume that the probability of the forward self-replication reaction (*P*_1→2_ or *P_SR_*) is much higher than the reverse reaction probability (*P*_2→1_ or 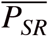), where the SRn process “undoes” itself following a time reversal trajectory. In fact, it has been suggested that the process of SRn is irreversible from a practical point of view (England, 2013; Saadat et al., 2020). Thus, the right hand side (RHS) term of eq. 1.1, 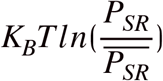, can be assumed to always have a positive value for a SR system undergoing SRn/growth (figure 1a). As shown in the original 2013 England paper the values for the terms in the ESL equation can be calculated with some certainty for simple reactions such as RNA and DNA replication and this approach provides valuable insights into the thermodynamic constraints controlling such reactions. However, calculating exact values for more complex reactions, such as the replication of a single bacterium, remained complex and required multiple assumptions that compromise the direct applicability of this analysis to even more complex systems (England, 2013).

**Figure 1:**
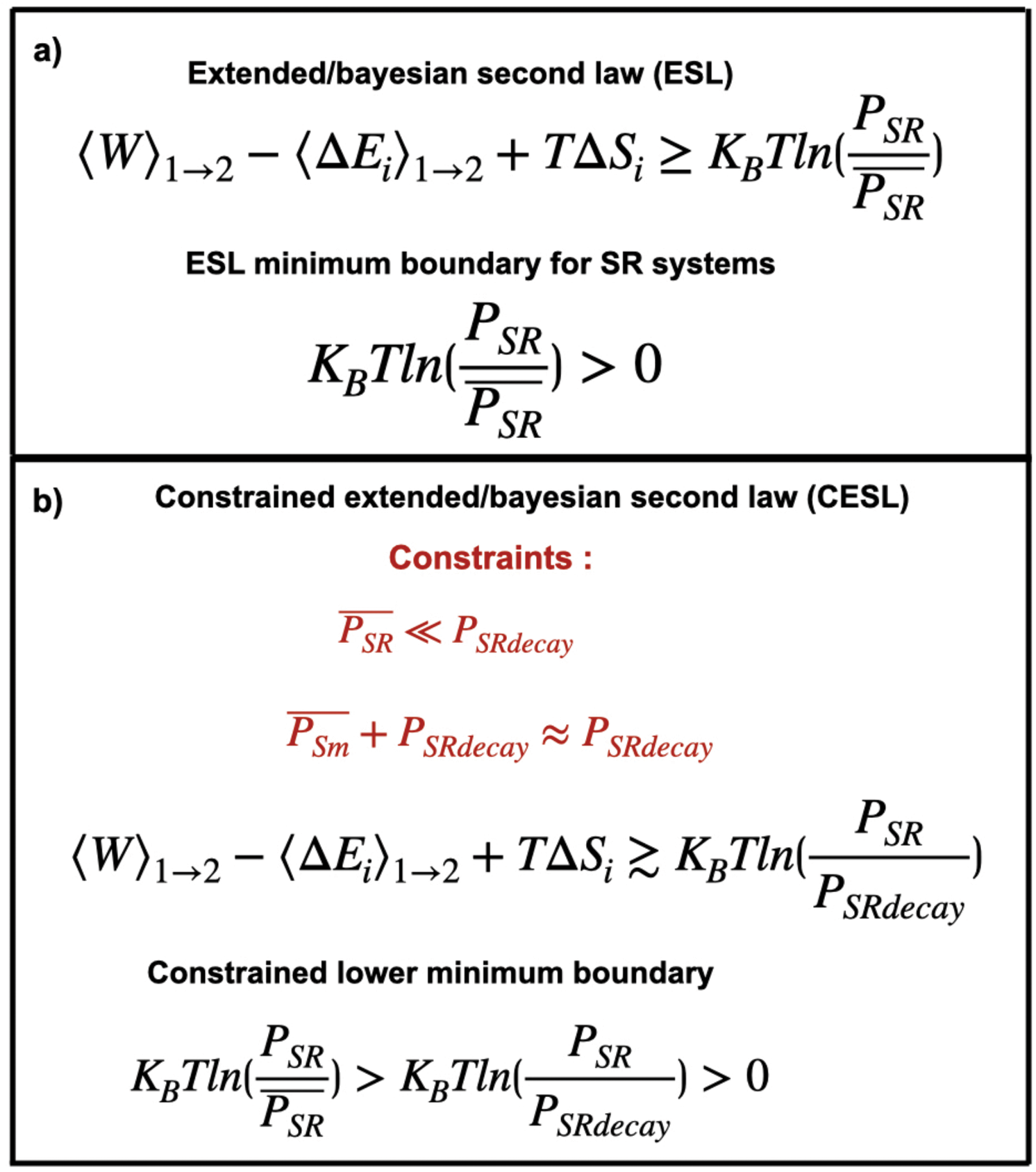
ESL a) and CESL b) equations with their associated minimum boundaries and constraints used in their derivation.

Nevertheless, in order to further take advantage of this analysis approach for more complex biological systems we can start by defining a simplistic model of SRn emergence, from a set of n reactants (*R_n_*) that combine themselves in a suitable fashion to generate a self-replicative entity *SR_m_* among a landscape of possible non-SR reaction products *A_m_* (abiogenesis phase, figure 2 top let panel). As addressed in the introduction, many unknowns still remain regarding the initial stage of the reaction where this SR entity emerges de novo. Fortunately, even without detailed knowledge of this initial reaction step/s we can extract some useful information from this model. Defining a bona fide SR system as inherently capable of growth (meaning specifically here: able to increase its overall complexity by increasing its internal energy and/or decreasing its internal entropy, (– *E_i_* + *S_i_*), concomitantly with an increase in biomass), when driven by an external field under suitable conditions allowing system growth and thus fulfilling the minimal requirements of life as discussed in the introduction, it then follows that the probability of SRn, *P_SR_*, must be intrinsically bigger than the SRn reverse probability plus the probability of system decay, 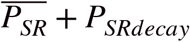, under this conditions. If we further assume that *P_SRdecay_* is substantially bigger than 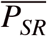, so that the likelihood of system decay is much greater than that of the system following the time reversal trajectory of SR, then we can simply assume that 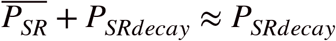, Introducing this into eq. 1.1 we obtain a Constrained Extended/Bayesian Second Law (CESL) equation (eq. 2; figure 1b and 2):

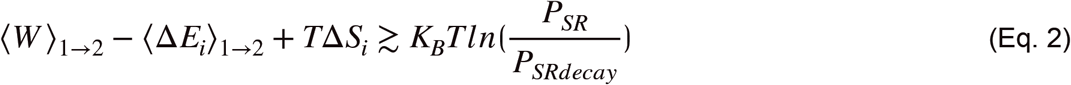

**Figure 2:**
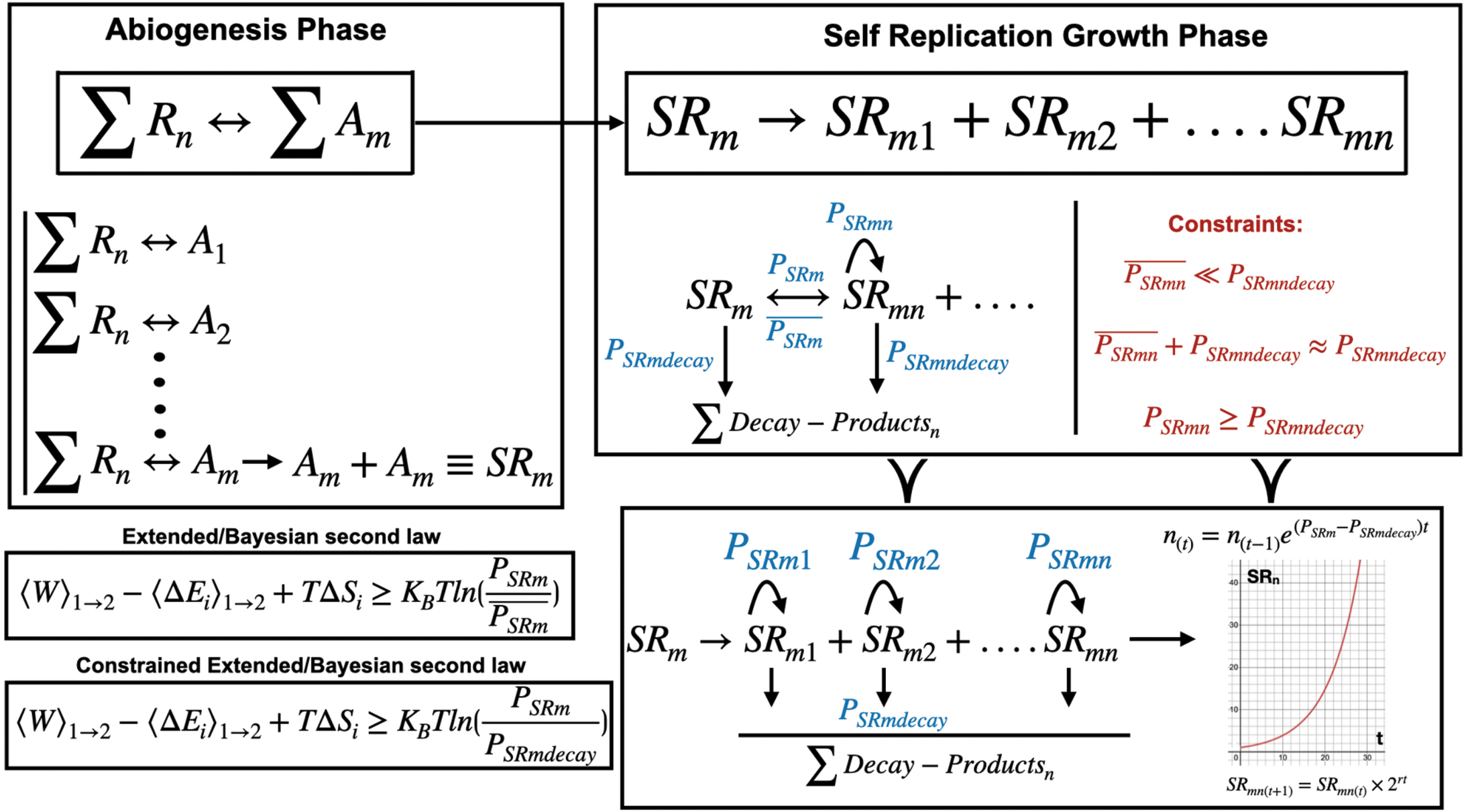
Model of original self replicator (SRm) emergence from non-self replicative reactants (R_n_) (abiogenesis) among the possible reaction assemblies (Am) and subsequent SR growth interpreted under the constraints set by the CESL equation, see text for further detail.

We can also easily see that for our system to grow *P_SR_* needs to be bigger than *P_SRdecay_* and therefore the RHS of Eq.2 would still be positive under this set of constraints, which resets a “new” constrained “lower” minimum boundary for the LHS of the ESL eq.: 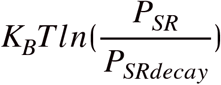 (Eq. 2 and figure 1b and 2 bottom left panel). This constrained equation is consistent with our observation of SR systems in nature at all levels, where system growth is observed whenever suitable conditions are present, from isolated bacterial growth (such as in a bio-incubator) (England, 2013; Saadat et al., 2020) to a mature complex ecosystem being perpetuated over time (Desmond-Le Quemener & Bouchez, 2014; Jorgensen, 2007; Kleerebezem & Van Loosdrecht, 2010; Kleidon, 2009).

From this initial constrained equation CESL it is then simple to see that under a suitable external driving field providing an unlimited supply of energy and given sufficient reactants, we would observe an exponential growing curve for our novel SR system over time modelled by an equation of the type *n*_(*t*)_ = *n*_(*t*–1)_*e*^(*P_SRm_-P_SRmdecay_*)*t*^, as previously described (see England, 2013 for details). This is again consistent with our experimental observation of equivalent life systems. For example, bacterial growth under a suitable environment progresses exponentially while resources (space, food, no competition, etc) are unlimited (figure 2, right hand side top and bottom panels: SR growth phase). Furthermore, bacterial specific theoretical thermodynamic growth models yield similar equations to the empirical Monod exponential growth equation, supporting the notion that thermodynamics underlies the growth behaviour of life systems as shown in figure 2, right hand side bottom panel, and modelled by an equation of the type *SR*_*mn*(*t*)_ = *SR*_*mn*(*t*–1)_ × 2^*rt*^ (Zeng & Yang, 2020), where r stands for time dependent bacterial growth constant and t for time.

It is worth to briefly acknowledge here the obvious observation that in complex biological systems the constrained probability term of the system, 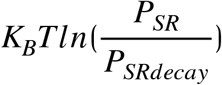, may not always be positive and lead to exponential growth as considered so far in our constrained system (figure 2, bottom right panel). Within the context of CESL, 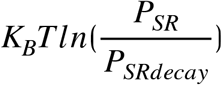 can take values (see figure 3):

- a positive value leading to exponential growth of the system and an increase in complexity due to an increase in internal energy and/or a decrease in internal entropy;
- a negative value leading to exponential decay of the system and a decrease in complexity due to a decrease in internal energy and/or an increase in internal entropy;
- be equal to 1, leading to a constrained driven “pseudo-equilibrium” where the available work is used in maintaining the system complexity (*E_i_* + *S_i_*) far from the thermodynamic equilibrium understood in the classical sense.

**Figure 3:**
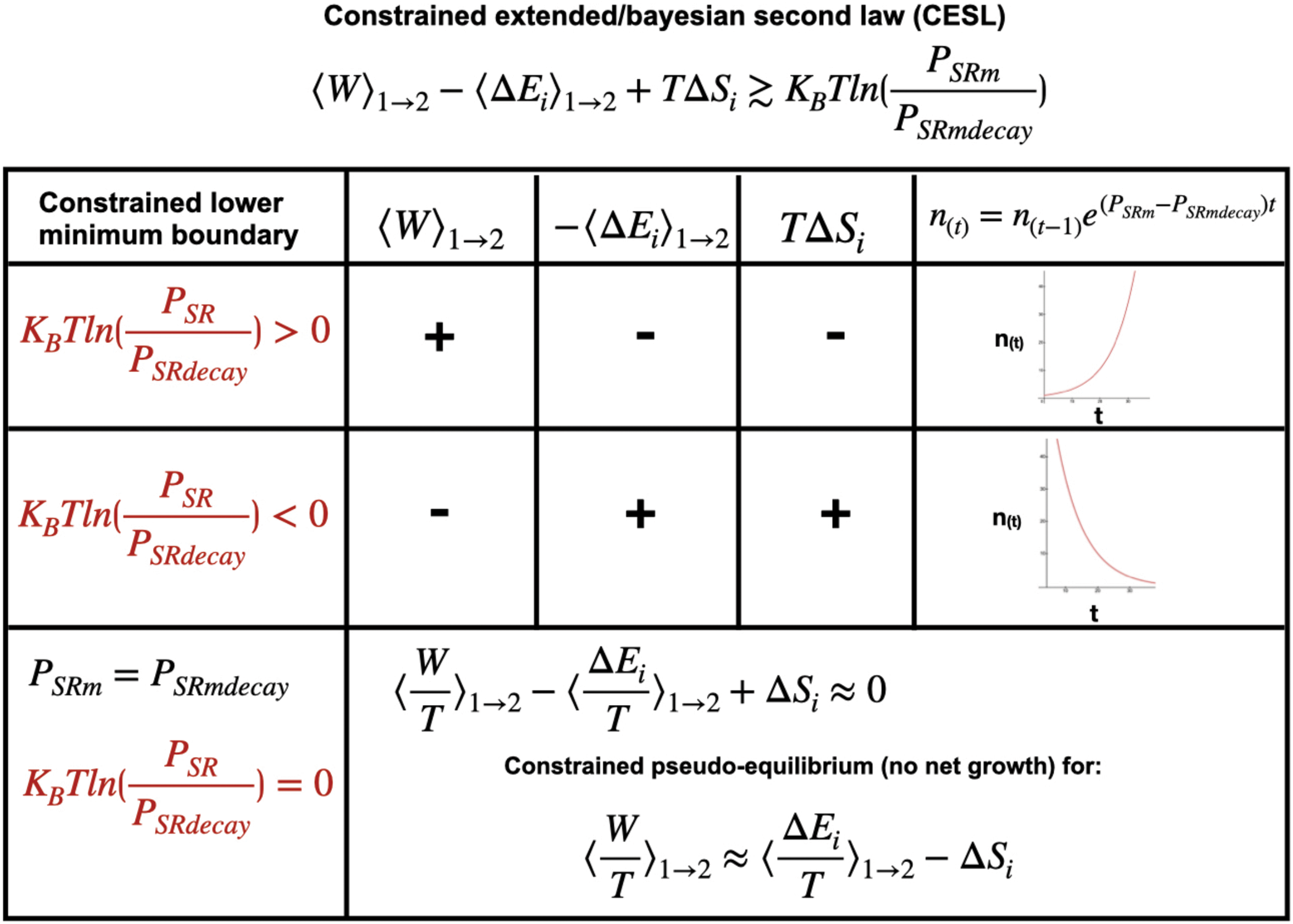
*Possible values of* 〈*W*〉_1→2_, 〈Δ*E_i_*〉_1→2_ *and T*Δ*S_i_ during growth, decay or pseudo-equilibrium evolution of a system with their associated minimum boundaries as constrained by the CESL equation.*

In particular we will focus in cases where the system undergoes growth or it is “held” in a driven pseudo-equilibrium where the available work is “used” in maintaining the system’s complexity (*E_i_* + *S_i_*) far from the thermodynamic equilibrium understood in the classical sense. In this constrained pseudo-equilibrium all usable work beyond the ESL minimum boundary level is used in maintaining the system’s complexity far from thermal equilibrium, with no net growth for *P_SR_* = *P_SRdecay_*, leading to: 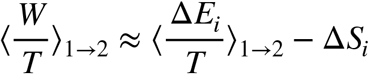 (see figure 3 bottom panel).

Importantly, in this context the system could still “self-renew” itself via SRn by transitioning from a CGS1 to a CGS2 that is equivalent in terms of overall complexity and biomass (*E_i_* + *S_i_*), such as a bacterial population grown inside a bioreactor where conditions are controlled in such a way that the total biomass is held constant while we have active cell division (because part of the bacterial population is harvested for a desired product for example). Furthermore, we propose that most stable mature complex ecosystem can be conceptually considered equivalent to the above system, self-renewing via SRn by transitioning from a CGS1 to a CGS2 that is equivalent in terms of overall complexity (and thus remaining stable).

Considering this further, we can also see that as far as the supply of usable work accessible to the system is unlimited and there are not intrinsic growth limiting factors that reduce 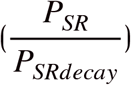(such as internal factors like durability vs decay or systemic factors such as competition or lack of resources that impede growth), and thus this term is a positive constant, the system would proceed to increase its internal energy and/or reduce its internal entropy exponentially according to the boundary limits set by eq. 2 under constrained conditions (and eq. 1 under general conditions). It is then self-evident that as long as a growing SR system is “loaded” with work from the drive it would exponentially increase its complexity (by reducing its internal entropy and/or increasing its internal energy and biomass) in a manner limited only by these inherent thermodynamic constraints, in the sense that there is direct correlation between the minimum boundary determined by the growth probability of the system and the work needed to increase its internal energy and decrease its entropy as described above and in England 2013; ie. the higher this lower minimum boundary (irreversibility) is the more work needed to to increase its internal energy and decrease its entropy in an equal amount.

In the SR systems analysed quantitatively in the original England 2013 paper (RNA, DNA and bacterial replication) it was assumed that the SRn process didn’t involved variation of the SR assemblies to allow for evolution as observed in nature. In order to shed further light on the behaviour of SR life systems at varying levels of complexity and evaluate system directionality over space and time we can re-examine this assumption by taking advantage of the capability to arbitrarily set the defining initial and final coarse grained system states in the ESL equation.

Keeping this goal in mind we can start by conceptually re-clarify that the two arbitrarily selected initial and final coarse grained states should include all individual reactants, products and SR assemblies needed for the system transition. For example, in the case of a single bacterium replication the initial coarse grained state would be composed of the bacterium plus all the reactants needed to facilitate SRn and the final state composed by the two resulting bacteria plus any generated products (figure 4 top panel). Importantly the theoretical justification for this interpretation is already described in detail in England, 2013 for this specific case. We propose that conceptually a similar interpretation can be made in a more complex system transition, such as the maturation process of an ecosystem from grassland to forest for example, where all the necessary reactants would be included into the initial coarse grained state together with all the individual SR entities present (the grassland ecosystem as a whole in this case: including bacteria, other decomposing organisms, plants, animals, etc) and the final state would also include any unused reactants, waste products and all the final SR entities present (the forest ecosystem as a whole including all components as before) (figure 4 bottom panel). Thus, for a far more complex system transition (ie. maturation process of an ecosystem from grassland to forest) we are still studying a closed system coupled to a water bath at constant volumes, temperature, etc and defining the initial and final coarse grained states to include all particles involved in the process (constant Ni) as required in ESL.

**Figure 4:**
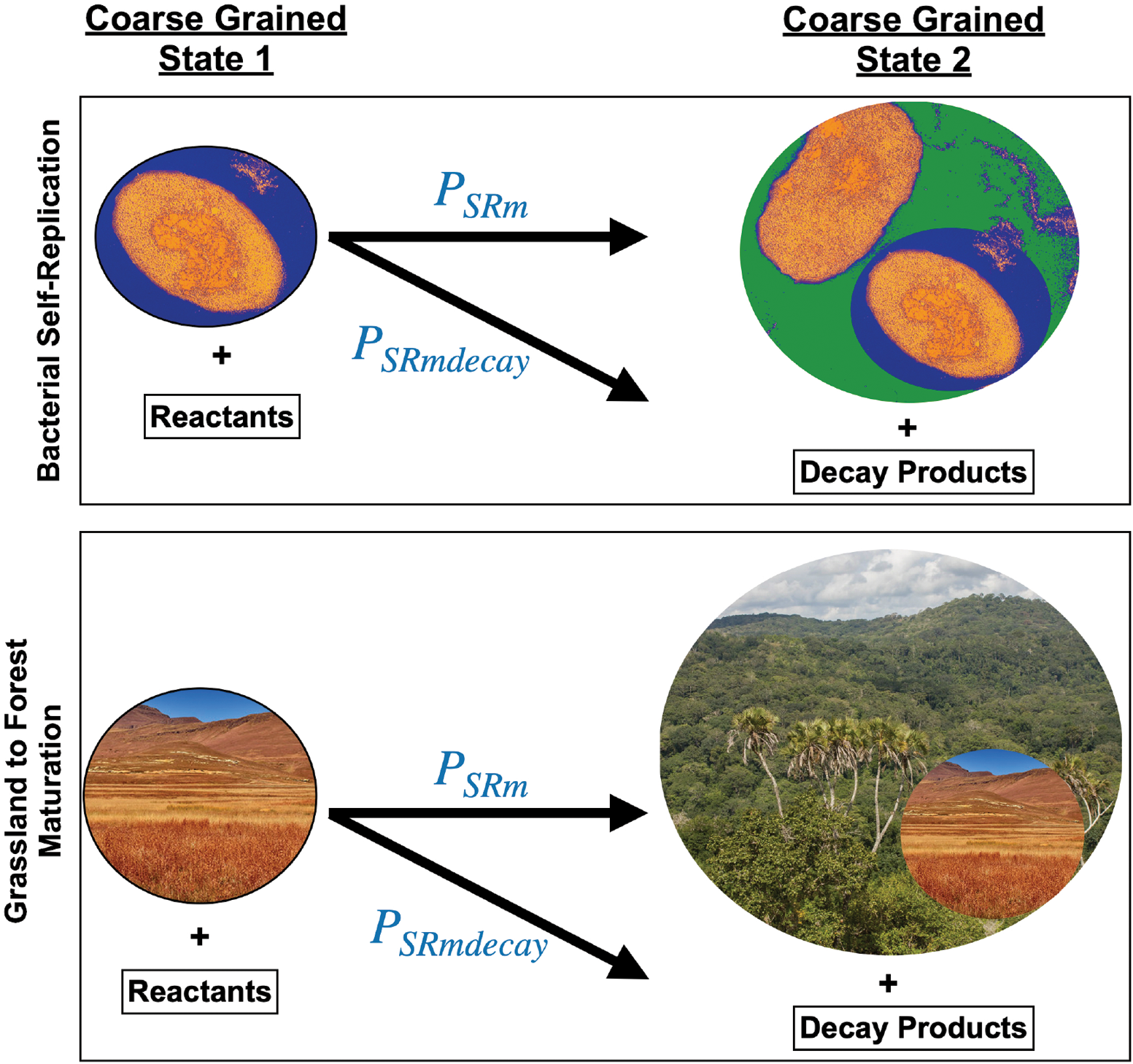
Model of the evolution of a complex life system over time, taking into account changes to “coarse graining” to allow for variation during the irreversible transition from a Coarse Grained State 1 to a Coarse Grained State 2 driven by SRn of the SR entities contained within the system at the different stages. As an example of two such systems the self-replication of a bacterium into two bacteria (top panel) and the self-replication (maturation) of a grassland into forest ecosystem (lower panel) are represented.

Importantly, however, the irreversibility of the self-replication of individual SR entities is still the “driving force” making the system transition irreversible as a whole as per our model described above (figure 2) and as described in England 2013 for a single bacterium SRn. We are simply expanding the “coarse graining” (ie grouping all individual SR entities as they share a equal properties in our analysis) of the initial and final states beyond the SRn reactions of individual SR entities within the system to include all particles within the system transition and thus still considering the coarse grained state transition irreversible as a whole. To put it another way, in England, 2013 bacterial SRn was studied using ESL by “coarse graining” all relevant metabolic reaction leading to SRn and defining CGS1 as 1 bacterium (plus reactants) and CGS2 as 2 bacteria (plus products) while ignoring the details of the intrinsically irreversible metabolic reactions leading to SRn. What we proposed here is exactly the same principle applied at a larger scale. We can apply “coarse graining” to individual SRn reactions (bacteria, grass, trees, etc) within the system in the same way that we applied it before to metabolic biochemical reactions (such as RNA and DNA replication needed for bacterial cell division) (England, 2013). In both cases the key aspect remains to define CGSs and system measurements compatible with the ESL model. Furthermore, the overall irreversibility restriction to the reaction is clearly as intuitively justified for the SRn of a single bacterium (England, 2013) as it is for the “SRn” (as a coarse grained state transition considered here) of a complex system, such as grassland maturing into forest ecosystem. If anything it could be argued that a forest ecosystem is even less likely to spontaneously revert to grassland following the time reverse trajectory due to its higher complexity (Jorgensen, 2007).

Finally, considering the system in this manner should also allows us to accommodate a small level variation of the individual SR entities (which could lead to adaptations and to the evolution of the overall SR system) within the constraints of the ESL model. With the hindsight of basic bacterial metabolism knowledge we can assume that two bacteria resulting from the SRn of one bacterium in a CGS1 to CGS2 transition as modelled by ESL would differ slightly in their intrinsic metabolic reactions. However, England 2013 has already demonstrated how to use the ESL model to study such and event and further showed that the theoretically calculated thermodynamic values for ESL eq. terms for bacterial duplication were very close to empirically measured values, validating the use of the ESL model for the study of bacterial replication. Following the same reasoning as above we can choose to apply a level of “coarse graining” such that allows for small changes to individual SRn reactions (bacteria, grass, trees, etc) within the system in the same way that we applied it before to metabolic biochemical reactions (such as DNA replication). In both cases the key still remains to define CGSs and system measurements compatible with the ESL model. We therefore propose that the initial and final coarse grained state SR individual entities may differ due to the progression of the system (in the above example during the transition from grassland to forest clearly a new layer of system complexity is added with different SR entities present) while still being able to apply the ESL model.

Therefore, the 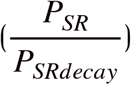 term would refer to the “replication” (CGS1 to CGS2 transition) of the whole complex system over time taking into account changes to the individual SR entities that allow for changes leading to evolution. Furthermore, as the system’s total particles must remain constant between CGS1 and CGS2, this allows us to evaluate changes in the extensive terms of CESL in a “semiquantitative” manner: a definition used here in the context of the ESL equation as the analysis of eq. terms that allows us to assign a directionality of change, or sign, to a specific term of interest within the equation given enough information (ie. at least the sign) regarding other terms is known.

Thus, we can now conceptually consider the evolution of the overall change in complexity of the system, specifically (-Δ*E_i_* + Δ*S_i_*) in ESL eq. In a biological system growing from a simpler coarse grained state 1 (CGS1) (i.e. grassland) to a more complex coarse grained state 2 (CGS2) (i.e. forest) we would expect this combined term to decrease (become more negative), as the system increases its *E_i_* by accumulating biomass and decreases its *S_i_* as it becomes more complex in terms of the number of component connections encompassed within the system. Importantly, it is already well established that on average biomass has a higher *E_i_* and lower *S_i_* than the inorganic reactants it comes from (Jorgensen, 2007). As all reactants (and products) are accounted within the CGS1 to CGS2 transition as described above we can calculate the directionality of change for the system’s (-Δ*E_i_* + Δ*S_i_*) from CGS1 to CGS2, even if we lack accurate numerical data. This approach has been intuitively used in the ecological sciences field (Jorgensen, 2007; Kleidon et al., 2010), but we can now describe it more formally in terms of thermodynamic equations. Thus this is a valuable approach as it allows us to study the term (−Δ*E_i_* + Δ*S_i_*) in the CESL model in a semiquantitative manner, ie assigning a directionality for its change based on a positive or negative change of the other terms of ESL eq. within the context of biological systems, even if we cannot calculate exact values for the eq. terms.

As we have described above (England, 2013), approximate values for the eq. terms can be calculated for simple reactions (such as RNA and DNA replication). However this is more difficult for more complex systems and even calculating the sign (directionality of change) for some terms in ESL eq. can be challenging. Therefore, it would be helpful to be further explore how to be able to empirically evaluate the individual terms of the CESL eq., at least in a semiquantitative manner that allows to explore the directionality of the reaction in more complex systems. To do so we can consider the following considerations regarding the individual ESL terms:

The term – 〈Δ*E_i_*〉_1→2_ represents the change in internal energy from a CGS1 to a CGS2. For biological systems we can simply approximate the internal energy of a system to the chemical energy resulting from the addition of the chemical potential of n components of the systems (Jorgensen, 2007; Wu et al., 2017). Even so it is still challenging to calculate an exact value for the internal energy of a complex system as it is technically difficult to measure the chemical potential of all its components. However, several directly proportional indicators have been used to calculate approximate values for a system’s internal energy (biomass, exergy, etc). By using these indicators we should able to assess the value of – 〈Δ*E_i_*〉_1→2_ in a semiquantitative manner that allows us to at least assign directionally to this change. We will focus hereafter on measurements of biomass, as these have been widely used and there is a wealth of data available. However, other directly proportional indicators such as exergy should provide equivalent results regarding the directionality of change in – 〈Δ*E_i_*〉_1→2_ (Fath et al., 2001). Biomass is clearly directly related to internal energy of a system resulting from its chemical potential energy and therefore we can assume that:

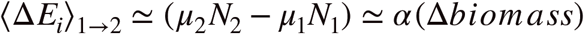

where *u*_1_ and *u*_2_ are the chemical potentials of the system, *N_i_* the number of chemical compounds and *α* a specific constant that relates the biomass of a system to its chemical potential energy. If we assume that *α* remains constant for CGS1 and CGS2 we can then assume that an increase in total biomass from CGS1 to CGS2 would result in a directly proportional increase in its internal energy. This is of course a crude approximation to real values as we are only evaluating chemical potential energy of the biomass in the system and we are omitting the chemical potential energy of inorganic compounds within the system (which should change in an inversely proportional manner as they are uptake to form biomass as the system grows). However, as biomass has an intrinsically higher chemical potential energy than its constituent inorganic compounds we can still reasonably assume that the change in biomass from CGS1 to CGS2 is directly proportional to the change in internal energy.

A similar analysis regarding the variation in internal entropy of the system, Δ*S_i_*, can also be put forward. The internal entropy term in ESL refers to the complexity of the system from a classical statistical mechanics point of view, thus relating the microstates probability distribution to the macroscopic system properties (in other words relating to the number of n components within the system and how these n elements are interconnected to one another). Measuring precisely the entropy changes in a complex system over time remains challenging. However, as above, we can evaluate changes in entropy in a semiquantitative manner that allows to at least explore the directionality of the reaction. For this we can take advantage of several ecological complexity orientators that are directly correlated to the internal entropy in a system such as ascendency, average mutual information (AMI), etc, and generally give a measure of how much energy flows are constrained within the system network and therefore reflecting the overall constrained network complexity as compared to the remaining degrees of freedom available to the system (Landauer, 1961; Ulanowicz, 2001). This approach has been used extensively within the network analysis field in ecology to evaluate system complexity and thus we can take advantage of previous experimental work. Importantly, recent work seeking to understand the complementarity of these orientators as ecological goal functions has shown that they are mutually consistent suggesting a common pattern in the network organisation of complex life systems (Fath et al., 2001). Thus measurement of Ascendency, AMI or even simply the number of total biological species within the system, would yield equivalent semi-quantitative results for the purpose of our analysis.Thus we can these measurements as a proxy indicator for the entropy evolution of a complex system, at least in a semiquantitative way as described above. As these indicators are inversely correlated with Δ*S_i_* an increase in Ascendency, AMI or total number off species from CGS1 to CGS2 would imply a decrease in *S_i_* (or increase in complexity) as described here.

Finally we can focus on the term reflecting the available work (energy) entering the system, 〈*W*〉_1→2_ from the external drive. Within the frame of the ESL model for the earth system we can assume that maximum work value limits are dependent of the capability of the system to input usable work from the external drive and this sets an upper limit for system growth and increase complexity (– Δ*E_i_* + Δ*S_i_*) for a given 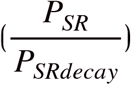 value. In other words, not all the energy available from the universe can enter the system due to its intrinsic limitations in terms of processing energy from the external drive. This can be clearly seen if we compared a non-photosynthetic system, where only chemical energy can be accessed as usable work, and a photosynthetic system, where a new energy level access directly from solar radiation can be used as work.

## Results

We are interested in extending the application of the ESL model to more complex biological systems than those already analysed in England, 2013 (RNA, DNA and single bacterium replication). We can begin by using data from complex ecological systems where detailed measurements of thermodynamic relevant parameters allows us to analyse changes in the ESL equation terms by selecting suitable coarse grained states to fit the available data.

In González et al., 2016, the authors study changes in the Tongoy Bay ecosystem using trophic network analysis. Tongoy Bay ecosystem was monitored over time from an initial CGS1 (1992), prior to which the ecosystem had suffered from anthropogenic over-exploitation, to a CGS2 (2012) where this anthropogenic disturbance had been ameliorated over time. The authors study changes in ecosystem structure and function using trophic network analysis to evaluate changes in macroscopic indexes related to the ecosystem’s health. Among these indexes we have data for biomass and ascendancy changes that we can input as described above into the ESL equation. Upon amelioration of the anthropogenic disturbance of the ecosystem, the authors describe an increase in the overall health, the total biomass and the interconnectivity (complexity) of the ecosystem by using several ecological indicators, showing that the system has experience a period of general recovery and growth from CGS1 to CGS2. Specifically, Ascendency (and therefore – *S_i_*) and biomass (and therefore *E_i_*) increased during this period (figure 5a) from 21127.2 to 28301.2 (flow bits), a 34.0 % increase, and 286.5 to 539.1 (gr m^-2^ year^-1^), a 88.2 % increase, respectively. We can also assume that 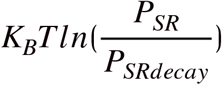 takes a positive value during the CGS1 to CGS2 transition, concomitant to a decrease in anthropogenic over-exploitation (this is reflected in reduced catch numbers on this period that would have resulted in an increased overall survival) (figure 5a, right panel), allowing for growth and establishing a positive thermodynamic minimum boundary according to ESL for the system. It then follows that the amount of work used by the system must have also increased in order to compensate for the increased in – *S_i_*, *E_i_* and positive 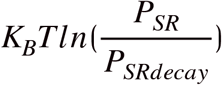, following the thermodynamic constraints set by ESL (figure 5a bottom panel). As no fundamental changes in the available usable work input into the system from the external drive are observed (such as the emergence of new properties like the photosynthesis example described above), we must conclude that this increase in work uptake must be inherent to the reported growth of SR entities capable of utilising work within it. This can be intuitively expected as higher biomass (88.2 % increase from CGS1 to CGS2) should be able to process more work from the external driving field. Thus, under the optics of ESL analysis we can conclude that the Tongoy Bay ecosystem had been kept under growth constraining conditions, with anthropogenic over-exploitation reducing 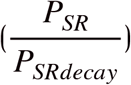, resulting in a driven pseudo-equilibrium (setting initial CGS1 conditions) during which the total amount of work available to the system was not fully utilised due to this growth limitation. Upon removal of this limitation (amelioration of anthropogenic over-exploitation), the system is then able to grow and uptake more work with a concomitant increase its overall complexity (Δ*E_i_* + Δ*S_i_*), until it “matures” and reaches a new state (CGS2) with a higher work uptake needed to maintain the system (figure 5a, bottom panel). Unfortunately, we do not have data regarding the evolution of albedo from CGS1 to CGS2, but assuming solar radiation as the main source of energy entering the system we would expect to see a concomitant reduction on overall albedo from CGS1 to CGS2.

**Figure 5:**
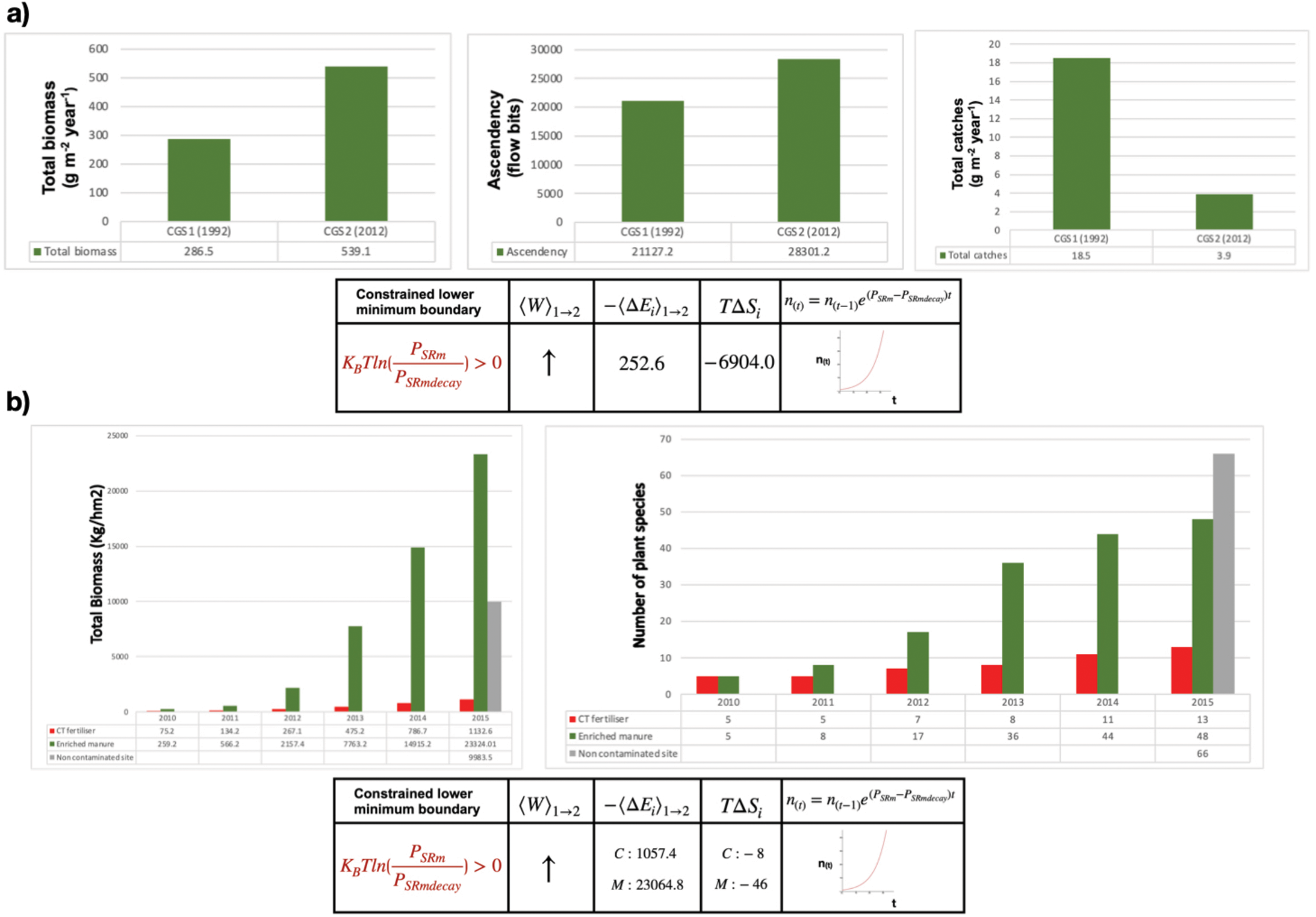
*Studying complex ecosystems by using trophic network analysis to evaluate changes in macroscopic indexes related to the ecosystem’s thermodynamic evolution as viewed under the ESL equation. In a) the evolution of the Tongoy Bay ecosystem was studied from CGS1 (year 1992) to CGS2 (year 2012) and changes in total biomass (top left graph), ascendency (top middle graph) and total catch (top right graph) were measured and used as directly proportional indicators for* 〈Δ*E_i_*〉_1→2_, *T*Δ*S_i_ and* 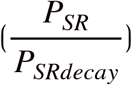 *respectively, within the terms of the ESL equation and the corresponding system progression (bottom panel), as justified in main text. In b) the evolution of the ecological restoration at a manganese mine contaminated wasteland ecosystem was studied from CGS1 (year 2010) to CGS2 (year 2015) and changes in total biomass (top left graph) and number of plant species (top right graph) were measured and used as directly proportional indicators for* 〈Δ*E_i_*〉_1→2_ *and T*Δ*S_i_ respectively, within the terms of the ESL equation and the corresponding system progression (bottom panel), as justified in main text above. A restoration strategy where experimental plots were modified by planting suitable wood plant species treated with enriched manure containing metal resistant bacteria was used (green columns). An equivalent control fertiliser only (red columns) and a control non-contaminated site (grey column) are included in the analysis.*

In a separate study a manganese mine contaminated wasteland that had undergone extensive plant biomass and species number depletion as compared to equivalent uncontaminated land was studied (Wu et al., 2017, Wu et al., 2018). Land at this contaminated wasteland showed high metal concentrations and low nitrogen and phosphorus levels. In an effort to establish a strategy for the ecological restoration of such ecosystems, experimental plots were modified by planting suitable wood plant species treated with enriched manure containing metal resistant bacteria or a chemical fertiliser only (containing equivalent levels of nitrogen, phosphorus and potassium) as a control. Plots were then monitored for 5 years (2010-2015). Total biomass, total number of plant species and total Gibbs free energy calculations for the plots were performed over the study period. These measurements showed a striking restoration of the ecosystem under the enriched manure treatment condition compared to the control. As above we can used ESL to gain insights into the changes in work and overall complexity in the system during this period using available data. As shown in figure 5b, left panel, the total biomass representing the overall *E_i_* of the system markedly increased over the experiment timeframe from 259.2 to 23324.0 (Kg hm^-2^), a 90 fold increase for the treatment; versus 75.2 to 1132.6 (Kg hm^-2^), a 15 fold increase for the control (with the equivalent Gibbs free energy measurement showing a similar trend) (figure 5b left panel and not shown). Similarly, there was a drastic decrease in the ecosystem entropy (*S_i_*), as shown by the increase in total plant species found compared to the chemical fertiliser internal control and approaching numbers similar to uncontaminated land: counted plant species increasing from 5 to 48 for treatment vs 5 to 11 for control, a 9.6 vs 2.2 fold increase respectively (figure 5b right panel); compared to 66 species counted in equivalent uncontaminated land. By artificially introducing new SR entities capable of growth within the contaminated system (wood plant species treated with enriched manure containing metal resistant bacteria) we have a situation similar from the point of view of ESL to the model shown in figure 2 relating to the emergence of the first SR entities during abiogenesis. We transition from a CGS1 with initial limited or null growth due to prior soil toxicity and poor nutrient availability limiting growth of pre-existing SR entities to a more complex CGS2 due to the artificial introduction in the system of novel SR entities capable of undergoing rapid growth thanks to their intrinsic properties: resistance to metals and available nutrients provided with the enriched manure (of note the artificially introduced wood plant species treated with enriched manure containing metal resistant bacteria must already be accounted for all parameters of interest in CGS1 in order to comply with ESL requirements). As expected these new SR entities drive growth of the overall system over time and lead to an increase in total *E_i_* and a decrease in *S_i_* which must come at the cost of a concomitant increase in the total work used by the system. The emergence of SR entities with new properties (resistance to metals and improved nutrient uptake) allows for an increased uptake of total usable work from the external drive. Thus, by reducing metal toxicity and providing additional nutrient resources that lead to a positive 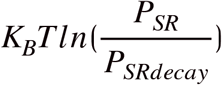 and allow an increase in *E_i_* we see an accompanying increase in – *S_i_* and usable work uptake inherent to the system’s growth (figure 5b, bottom panel).

Interestingly, in an somewhat controlled experimental set-up we can observe under the light of ESL that the system progresses to higher levels of complexity as changes within its structure allow it to further exploit improved growth conditions in a positive feed-back loop (further reduction of toxic metal in the soil and enrichment of nutrient availability allowing other SR entities to grow), driven by the initial emergence (controlled introduction in this context) of the novel SR entities, in an manner that seems analogous to the process of complex ecological systems evolution during relevant historical periods as conceptually described in the Gaia hypothesis (Karnani et al., 2009).

In order to further explore this idea we can design a thought experiment where we evaluate the thermodynamic progression of a hypothetical SR biological system from evolutionary relevant coarse grained state 1 (CGS1) to 2 (CGS2) in terms of the ESL equation.

We can set our CGS1 and CGS2 at evolutionary relevant states for SR systems during early earth history, and thus we can study how changes in usable work (*W_usable_*) levels relate to changes in complexity, (–Δ*E_i_* + Δ*S_i_*), and growth, 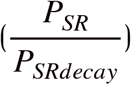, over evolutionary relevant time frames.

First let us assume that in our SR system usable work would be accessed overall from anaerobic chemical reactions, anaerobic photosynthesis, aerobic photosynthesis and aerobic chemical reactions with known values of energy available for each of these reaction types based on generic empirical data for commonly used substrates/reactions in bacterial metabolism (figure 6a). We can define a CGS1 where usable work only enters the system via anaerobic bacteria capable of anaerobic chemical reactions (with an overall value of total energy availability derived from all commonly used anaerobic substrates described in 6a combined). This results in a total amount of usable work at CGS of level 1, or *W*_*usable*_1__, under anaerobic conditions such as those present on earth until approximately 2.5 billion years ago with low oxygen levels (*pO*_2_ < 0.1 % *PAL*) (PAL: present atmospheric level) (anaerobic metabolism in figure 6b for combined total system value and 6c for individual contributing reactions). Let us further assume that our SR system in CGS1 undergoes intrinsic changes during the conversion to CGS2 in an evolutionary relevant time frame resulting on the emergence of novel properties allowing the system to access energy directly from sunlight due to the appearance of first anaerobic and second aerobic photosynthesis and leading to a global oxygenation event (GOE) and a rise in oxygen levels to at least *pO*_2_: 1 – 10 % *PAL.* Thus setting conditions such as those present on earth from approximately 2.5 to 0.5 billion years ago, which in turn allows for aerobic chemical reactions and the emergence of aerobic bacteria and results in a system with anaerobic metabolism, anaerobic photosynthesis, aerobic photosynthesis and aerobic metabolism present (6b top panel for combined total system values and 6c for individual contributing reactions within the proposed timeframe). This final state corresponds to CGS2. We can now observe that our CGSs transition is organised in a hierarchical configuration of increasing complexity levels corresponding to the historical development of the system. Thus, CGS2 includes all previous relevant modes of accessing work (anaerobic chemical reactions) plus all emergent modes developed during the transition (anaerobic photosynthesis, aerobic photosynthesis and aerobic chemical reactions),which in combination define the usable work at CGS level 2, or *W*_*usable*_2__ (figure 6b top panel, all combined). Based on the available energy from these modes of accessing work we can calculate total *W_usable_* per reaction (kJ per mol of electron donor) at CGS level 1 and 2 (figure 6a and 6b top panel). As expected, *W*_*usable*_2__ > *W*_*usable*_1__ (743.2 vs 9345.2 Kj per mol electron donor respectively), in this case by more than ten fold (12.57 fold increase), as CGS2 includes the available work at CGS1 (anaerobic chemical reactions) plus the work accessible by the new work modes (anaerobic photosynthesis, aerobic photosynthesis and aerobic chemical reactions) (figure 6a, 6b top panel and 6c) (Heijn et al., 1992; Roden et al., 2011; McCarty, 2007).

**Figure 6:**
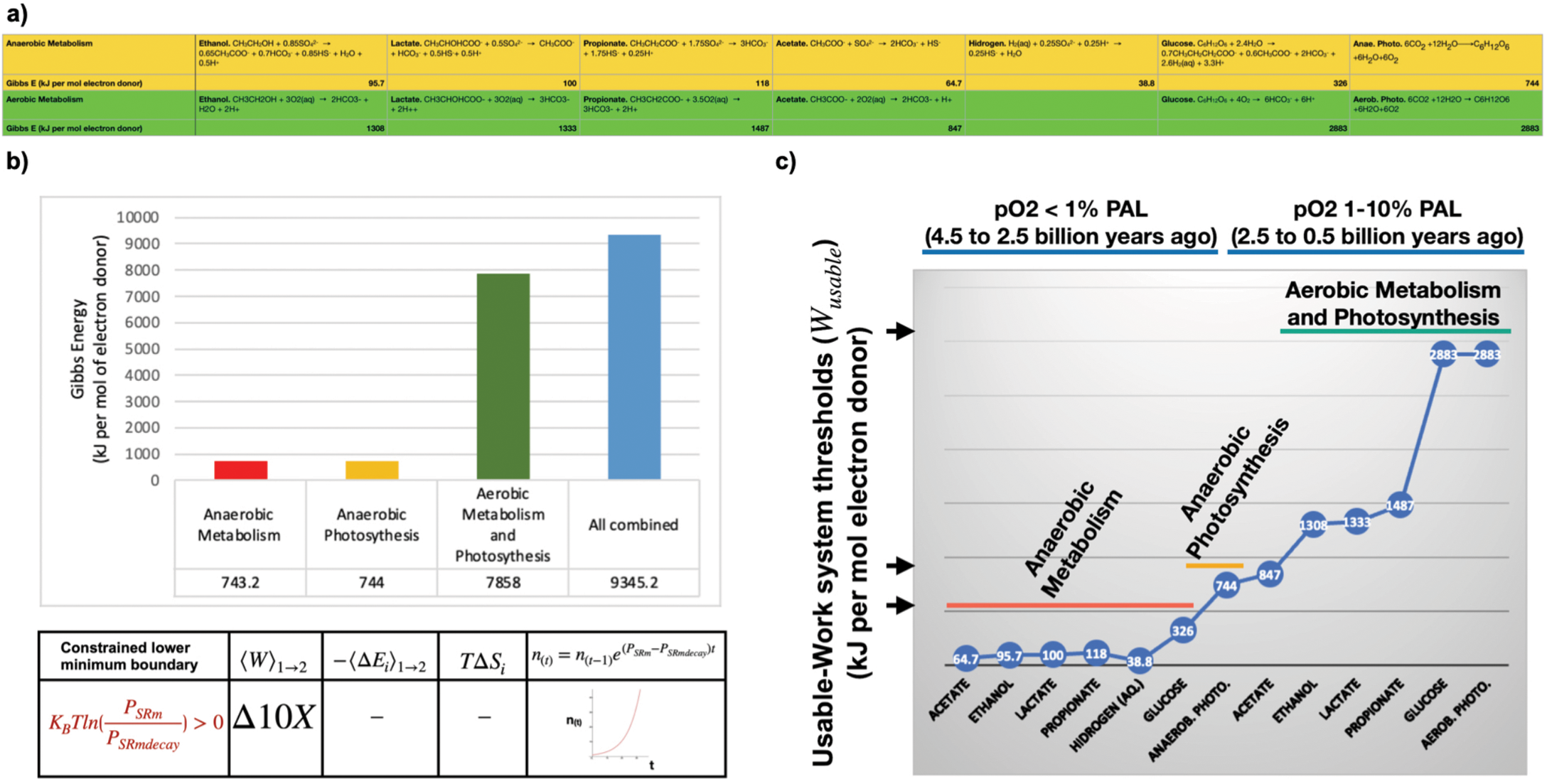
A thought experiment where we analyse the hypothetical thermodynamic progression in terms of the ESL equation of a complex prokaryotic biological system from evolutionary relevant to earth’s early life history CGS1 to CGS2. In a) the free energy (kJ per mol of electron donor) of evolutionary relevant prokaryotic aerobic, anaerobic (respiratory and/or fermentative) metabolism of glucose, ethanol, formate, acetate, lactate, propionate and H_2_ and aerobic and anaerobic photosynthesis reactions are shown. In b) the free energy values for combined anaerobic and/or aerobic reactions using data form a) as described is calculated. For example, the anaerobic metabolism column in red adds the value all the possible non-photosynthetic anaerobic reactions included in a) as a representation of the possible relevant anaerobic modes of energy access. Combined values for anaerobic photosynthesis in yellow, aerobic metabolism and photosynthesis in green and total combined values in blue are calculated equivalently. In c) changes in usable-work system thresholds (i.e. total amount of W_usable_ that can be accessed via the free energy generated from anaerobic and/or aerobic reactions) relating to prokaryotic evolution and correlated levels of atmospheric oxygen during early history of life on earth are represented.

We have thus moved from *W*_*usable*_1__ to *W*_*usable*_2__ following a relevant evolutionary time frame of events (figure 6c) (Catling et al., 2005; Sessions et al., 2009; Soo et al., 2017; Yamamoto et al., 2011). We do not know the total exact values for *W*_*usable*_1__ and *W*_*usable*_2__, as we lack reliable historical data regarding exact biomass changes from CGS1 to CGS2. However, it is safe to assume that with the advent of photosynthesis organisms would have colonised new ecosystems and the overall biomass would have been greatly increased during the equivalent evolutionary historical period (from only anaerobic to anaerobic+photosynthetic+aerobic prokaryotic ecosystems). Importantly, calculations based on historical records are in agreement with this assumption (Battistuzzi et al., 2004; Catling et al., 2005; Payne et al., 2009).

Therefore, we can safely assume that:

1. *W*_*usable*_2__ > *W*_*usable*_1__, to which data from figure 6b and 6c is an underestimation as it only relates to intrinsic access to usable work per reaction type, but it doesn’t account for total biomass increases, and thus 〈*W*〉_1→2_ > 0 (figure 6b bottom panel).
2. We can also safely assume that during this evolutionary relevant period 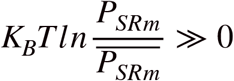 and 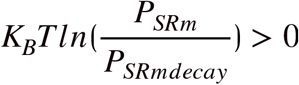, as we have multiple evidence of new organism types colonising newly available ecosystems due the novel evolutionary solutions that appeared in this time with an inherent positive growth to our system (Catling et al., 2005; Hamilton et al., 2016; Hsia et al., 2013; Payne et al., 2009) (figure 6b bottom panel).

It then follows that an increase in the total complexity of the system (–Δ*E_i_* + Δ*S_i_*) from CGS1 to CGS2 would be expected under the constraints imposed by ESL (figure 6b bottom panel). This is in agreement with empirical observations of this period. As already explained above, internal energy should increase in line with the expected increase in total biomass from CGS1 to CGS2. Likewise, entropy should decrease accordingly with the complexity of the system as more levels are added to its hierarchical structure. This appears self-evident from the evolutionary history followed in hearth during the period that we have chosen to model. In CGS1 we only have anaerobic prokaryotes ecosystems, limited to local chemical gradients that allow for anaerobic chemical reactions maintaining the system. These could be loosely modelled after modern geothermal vent ecosystems (Yamamoto et al., 2011) (figure 7). A CGS1 pseudo-equilibrium state with a maximum work value limit dependent of the capability of the system to input usable work from the external drive (anaerobic chemical reactions) sets an upper limit for system growth and increase complexity according to ESL (figure 7). During this initial pseudo-equilibrium state with no net growth we have 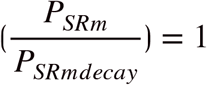 and CGS1 cycles itself in a maintenance loop with 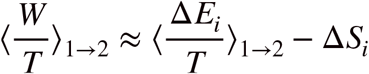 and thus most of the usable work is used in maintaining the system overall complexity (–Δ*E_i_* + Δ*S_i_*) far from equilibrium as understood in the classical sense. In order for the system to undergo changes in complexity it must be able to access more usable work from the universe. This can be achieved via the development of novel intrinsic adaptations that allow to access a higher level of work, such as being able to perform photosynthesis or aerobic respiration (figure 7) and transition from CGS1 to CGS2 as described above. Of note here the individual SR entities (prokaryotes in this model) may not change their intrinsic complexity substantially themselves (an anaerobic prokaryote vs an evolutionary related aerobic photosynthetic prokaryote), however the complexity reflected in ESL refers to the overall system in a hierarchical manner, which clearly increases its complexity greatly in both terms its internal energy and its internal entropy (–Δ*E_i_* + Δ*S_i_*), and comparing the complexity of locally isolated anaerobic bacterial communities vs aerobic photosynthetic communities widespread throughout the planetary surface we can immediately appreciate the scale of overall system change (figure 7) (Battistuzzi et al., 2004; Hamilton et al., 2016; Hsia et al., 2013; Payne et al., 2009). Thus, during the CGS1 to CGS2 transition the system can input a higher level of usable work that can be used in a phase of exponential growth to offset the cost of its increase complexity according to the minimum boundary constraints imposed by ESL (figure 7). Without further data regarding the evolution of the system after CGS2 we can assume that the CGS1 to 2 transition ends when a CGS2 pseudo-equilibrium is reached and the system would then cycle itself in a maintenance loop with a higher work input requirement than that of the CGS1 pseudo-equilibrium due to its overall higher complexity, as required by ESL (figure 7). In time the emergence of novel evolutionary relevant properties allowing access to novel modes of accessing work (such as the emergence of eukaryotic cells and multicellularity) could lead to an further cycle of hierarchical complexity expansion and a reset of the minimum boundary constraints imposed by ESL (figure 7).

**Figure 7:**
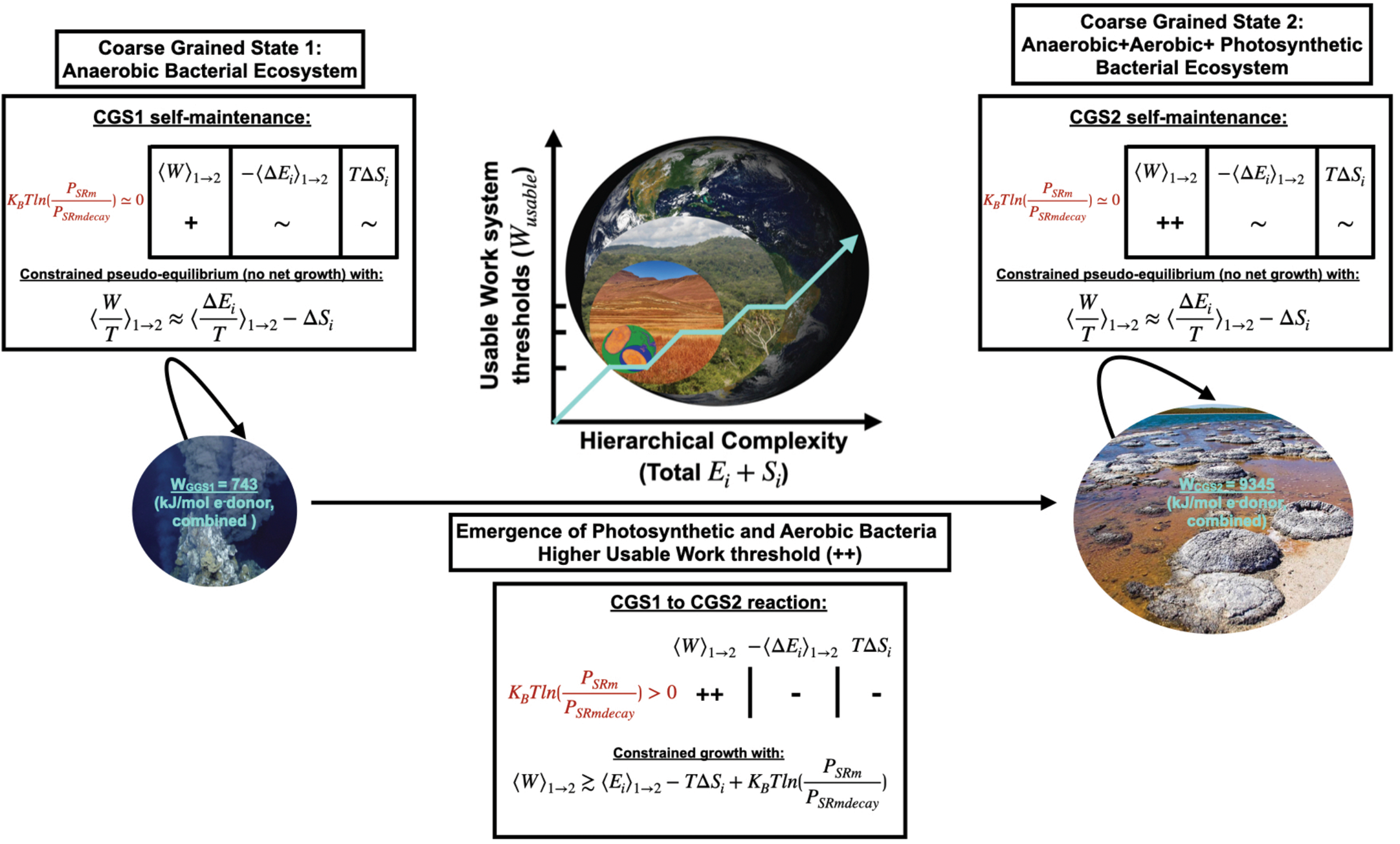
A complete model of the thermodynamic progression of a complex prokaryotic biological system from evolutionary relevant CGS1 (anaerobic) to CGS2 (aerobic and anaerobic). The system would adopt a thermodynamic hierarchical organisation with increasing levels of complexity over the modelled evolution time as higher levels of energy (higher usable work thresholds) are accessed, according to minimum boundary constraints set by the ESL equation.

## Discussion

The ESL thermodynamic equation allows us to study far from equilibrium driven self-replicative life systems undergoing irreversible coarse grained state transitions. Meaningful approximate values for the ESL equation terms can be calculated for simple SR systems (England, 2013) and offer important insights into the behaviour such systems. We have further proposed that ESL based semi-quantitative analysis of more complex SR systems (for which calculating values for the ESL equation terms is more challenging) also offers valuable insights into the progression of such systems in the context of the thermodynamic constraints underlining system behaviour.

Specifically, we have seen how using data from complex ecosystem modelling we can apply the ESL equation to interpret in a semi-quantitative manner the thermodynamic changes that underpin the system’s evolution. Likewise, we have shown how this analysis can be extended to interpret relevant evolutionary ecological timeframes of life on earth in a semi-quantitative manner, even in the absence of accurate experimental data. These results agree with current experimental observations (González et al., 2016; Holdaway et al., 2010; Schneider et al.,1994; Wu et al., 2017; Wu et al., 2018) and with our intuitive understanding of the organisation and evolution of life systems on earth (Jorgensen, 2007; Kleidon, 2009; Prigogine & Nicolis, 1971) and lead us to propose that self-replicative life systems are evolutionary and spatially organised in hierarchical thermodynamic levels of incremental work usage inputted from the external driving field (figure 7). In this context for a SR system to be able to overcome its upper bound complexity threshold set by ESL it must then be able to access higher levels of usable work/energy from the universe, as defined by the minimum boundaries set by ESL for irreversible SR systems (figure 1 and 3). This can be achieved through hierarchical variation in the properties of the system in a relevant evolutionary timescale and multiple examples of this emergence of new levels of usable work underpinned the evolution of life on hearth (such as in the evolution of photosynthetic SR organisms exemplified above). Furthermore, the more work that can be inputted into the system from the universe the lower the unused or reflected work would be, and this can be seen as a lower albedo as complexity increases in life systems. The example of grassland vs forest ecosystem is appropriate here again, with a more complex forest ecosystem generating a lower albedo that a grassland ecosystem, which in turn also generates a lower albedo than an even simpler dessert ecosystem (Holdaway et al., 2010; Schneider et al., 1994).

This ESL based interpretation provides valuable insights regarding the initial emergence of SR systems as well. As we have seen this approach supports that, in the presence of an external driving field and under suitable conditions allowing growth, SR entities would self-replicate growing exponentially according to the constraints set by Eq. 1.1 (figure 2). It has already been hypothesised that, from a theoretical thermodynamics analysis point of view, biological systems are constrained to evolve favouring selection that leads to configurations that maximise positive values of work; in other words, evolution leading to arrangements of the system that are more capable of inputting the work available to them from the external drive (England, 2015; Prigogine & Nicolis, 1971). Importantly, this approach allows us to explore this hypothesis by arbitrarily setting the initial and final coarse grained states to study. Thus, we can define the initial state (CGS1) as the set of n reactants and the final state (CGS2) as the emergent m assemblies, among which systems capable of SR would emerge during abiogenesis (figure 2). Many trajectories are theoretically possible from the initial set of reactants leading to emergent systems capable of SR. However, maintaining entropy changes constant (ie: the novel m emergent assemblies are all of equivalent complexity) and under non limiting resources conditions, assembly configurations more capable of maximising positive values of work, (ie more capable of using the work driven into the system from the external drive) would be able to maximise their exponential growth (maximise 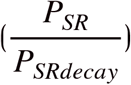) (figure 2 and eq. 3) (England, 2015; Perunov et al., 2016) and quickly dominate the configuration landscape. As during abiogenesis the initial de novo emergent SR systems are unlikely to have encountered strong competition environments and/or limiting resources this analysis may provide an approximation to the conditions under which life emerged on earth. Conceptually, similar experiments studying early exponential growth stages of easily monitored de novo system, such as inoculation of one bacterium in a suitable growth media or the clonal selection of rapidly growing tumour cells from a heterogeneous population seem to support this hypothesis (Greaves et al., 2012). However, it is challenging to faithfully replicate these conditions artificially and thus further work remains needed to validate this hypothesis experimentally.

From a general empirical point of view it could be argued that the fundamental postulates of statistical mechanics could be reinterpreted in reverse for far from equilibrium self-replicative life systems driven by an external driving field. We propose that under the optics of the ESL model the external driving field imposes work-dissipation evolutionary constraints on the system with boundaries set by the total usable work available to the system, or in other words, the work that can access or be loaded into the system’s assemblies due to their intrinsic molecular arrangements. Thus, under this constraint the system evolves to adopt higher work dissipation macrostates within the system’s permissible microstate configurations, and this is reflected in a decrease of the total entropy in the system (–Δ*E_i_* + Δ*S*). As self-replicative growth and variation are intrinsic features of bona fide life SR systems (as long as the external drive inputs/loads work into the system), this constraint would drive the evolution of the system to follow an increase in its complexity path under stable growth conditions. Therefore, in our proposed hierarchical thermodynamic interpretation, the work from the external driving field loaded into the system can be viewed as an evolutionary pressure selecting for higher work dissipative system arrangements. These in turn will further follow further evolutionary trajectories that maximise positive values of work and thus constraining the system to evolve to higher levels of work dissipation/use as its intrinsic features change over evolutionary relevant time frames. Importantly, this prediction based on thermodynamics analysis agrees with all our empirical observations of the evolution of self-replicative life systems on earth (England, 2015; Jorgensen, 2007; Kleidon, 2009; Perunov et al., 2016; Schneider et al.,1994), implying that the basic organisation of self-replicative life systems is constrained by these basic thermodynamic principles. Thus, Fisher’s remark “Natural selection is a mechanism for generating an exceedingly high degree of improbability” could be flipped upside down and reinterpreted as: “Generating an exceedingly high degree of improbability is an inevitable consequence of a hierarchical thermodynamic natural selection mechanism”.

Finally, we have shown that from a practical point of view complex ecosystems (such as the restoration effect on a manganese tailing wasteland ecosystem seen in results) and relevant evolutionary ecological time frames of life on earth can be interpreted using a thermodynamic semiquantitative analysis. Interestingly we can see how punctuated evolutionary stages of life on earth correlate to incremental levels of work loaded into the system by the external drive with a corresponding increase in system’s complexity (figure 6 and 7). We proposed a thermodynamic interpretation for the clear trend towards global increased complexity of the earth-life system, in agreement with current theories (Karnani et al., 2009; Walker, 2017), and proposed that in the absence of external perturbations the directionality of this path (increased complexity and work dissipation) is set by the thermodynamic evolutionary constraints discussed in this manuscript. We propose that this type of analysis would be useful in understanding the directionality of complex biological systems across time and space and that experimental validations using artificially designed models could be easily implemented to further corroborate its validity.

## References

Attwater, J., Wochner, A., Pinheiro, V. B., Coulson, A., & Holliger, P. (2010). Ice as a protocellular medium for RNA replication. Nature Communications, 1. https://doi.org/ARTN7610.1038/ncomms1076

Battistuzzi, F. U., Feijao, A., & Hedges, S. B. (2004). A genomic timescale of prokaryote evolution: insights into the origin of methanogenesis, phototrophy, and the colonization of land. Bmc Evolutionary Biology, 4. https://doi.org/Artn4410.1186/1471-2148-4-44

Bartolotta, A., Carroll, S. M., Leichenauer, S., & Pollack, J. (2016). Bayesian second law of thermodynamics. Phys Rev E, 94(2-1), 022102. https://doi.org/10.1103/PhysRevE.94.022102

Catling, D. C., Glein, C. R., Zahnle, K. J., & Mckay, C. P. (2005). Why O-2 is required by complex life on habitable planets and the concept of planetary “oxygenation time”. Astrobiology, 5(3), 415–438. https://doi.org/DOI10.1089/ast.2005.5.415

Crooks, G. E. (1999). Entropy production fluctuation theorem and the nonequilibrium work relation for free energy differences. Physical Review E, 60(3), 2721–2726. https://doi.org/DOI10.1103/PhysRevE.60.2721

Desmond-Le Quemener, E., & Bouchez, T. (2014). A thermodynamic theory of microbial growth. Isme Journal, 8(8), 1747–1751. https://doi.org/10.1038/ismej.2014.7

Duim, H., & Otto, S. (2017). Towards open-ended evolution in self-replicating molecular systems. Beilstein J Org Chem, 13, 1189–1203. https://doi.org/10.3762/bjoc.13.118

England, J. L. (2013). Statistical physics of self-replication. J Chem Phys, 139(12), 121923. https://doi.org/10.1063/1.4818538

England, J. L. (2015). Dissipative adaptation in driven self-assembly. Nat Nanotechnol, 10(11), 919–923. https://doi.org/10.1038/nnano.2015.250

Fath, B. D., Patten, B. C., & Choi, J. S. (2001). Complementarity of ecological goal functions. J Theor Biol, 208(4), 493–506. https://doi.org/10.1006/jtbi.2000.2234

Gonzalez, J., Ortiz, M., Rodriguez-Zaragoza, F., & Ulanowicz, R. E. (2016). Assessment of longterm changes of ecosystem indexes in Tongoy Bay (SE Pacific coast): Based on trophic network analysis. Ecological Indicators, 69, 390–399. https://doi.org/10.1016/j.ecolind.2016.04.019

Greaves, M., & Maley, C. C. (2012). Clonal evolution in cancer. Nature, 481(7381), 306–313. https://doi.org/10.1038/nature10762

Hamilton, T. L., Bryant, D. A., & Macalady, J. L. (2016). The role of biology in planetary evolution: cyanobacterial primary production in low-oxygen Proterozoic oceans. Environmental Microbiology, 18(2), 325–340. https://doi.org/10.1111/1462-2920.13118

Heijnen, J. J., & Van Dijken, J. P. (1992). In search of a thermodynamic description of biomass yields for the chemotrophic growth of microorganisms. Biotechnol Bioeng, 39(8), 833–858. https://doi.org/10.1002/bit.260390806

Holdaway, R. J., Sparrow, A. D., & Coomes, D. A. (2010). Trends in entropy production during ecosystem development in the Amazon Basin. Philosophical Transactions of the Royal Society B-Biological Sciences, 365(1545), 1437–1447. https://doi.org/10.1098/rstb.2009.0298

Hsia, C. C. W., Schmitz, A., Lambertz, M., Perry, S. F., & Maina, J. N. (2013). Evolution of Air Breathing: Oxygen Homeostasis and the Transitions from Water to Land and Sky. Comprehensive Physiology, 3(2), 849–915. https://doi.org/10.1002/cphy.c120003

Jorgensen, S. E. (2007). A new ecology: systems perspective. Elsevier. http://www.ezproxy.is.ed.ac.uk/login?url=https://www.sciencedirect.com/science/book/9780444531605

Jarzynski, C. (2011). Equalities and Inequalities: Irreversibility and the Second Law of Thermodynamics at the Nanoscale. Annual Review of Condensed Matter Physics, Vol 2, 2, 329–351. http://www.ezproxy.is.ed.ac.uk/login?url=https://www.sciencedirect.com/science/book/9780444531605

Jarzynski, C. (2011). Equalities and Inequalities: Irreversibility and the Second Law of Thermodynamics at the Nanoscale. Annual Review of Condensed Matter Physics, Vol 2, 2, 329–351. https://doi.org/10.1146/annurev-conmatphys-062910-140506

Jorgensen, S. E., Ladegaard, N., Debeljak, M., & Marques, J. C. (2005). Calculations of exergy for organisms. Ecological Modelling, 185(2-4), 165–175. https://doi.org/10.1016/j.ecolmodel.2004.11.020

Karnani, M., & Annila, A. (2009). Gaia again. Biosystems, 95(1), 82–87. https://doi.org/10.1016/j.biosystems.2008.07.003

Kleerebezem, R., & Van Loosdrecht, M. C. M. (2010). A Generalized Method for Thermodynamic State Analysis of Environmental Systems. Critical Reviews in Environmental Science and Technology, 40(1), 1–54. https://doi.org/Pii91837523310.1080/10643380802000974

Kleidon, A. (2009). Nonequilibrium thermodynamics and maximum entropy production in the Earth system. Naturwissenschaften, 96(6), 653–677. https://doi.org/10.1007/s00114-009-0509-x

Kleidon, A., Malhi, Y., & Cox, P. M. (2010). Maximum entropy production in environmental and ecological systems. Philos Trans R Soc Lond B Biol Sci, 365(1545), 1297–1302. https://doi.org/10.1098/rstb.2010.0018

Landauer, R. (1961). Irreversibility and Heat Generation in the Computing Process. Ibm Journal of Research and Development, 5(3), 183–191. https://doi.org/DOI10.1147/rd.53.0183

McCarty, P. L. (2007). Thermodynamic electron equivalents model for bacterial yield prediction: modifications and comparative evaluations. Biotechnol Bioeng, 97(2), 377–388. https://doi.org/10.1002/bit.21250

Meysman, F. J. R., & Bruers, S. (2010). Ecosystem functioning and maximum entropy production: a quantitative test of hypotheses. Philosophical Transactions of the Royal Society B-Biological Sciences, 365(1545), 1405–1416. https://doi.org/10.1098/rstb.2009.0300

Nakashima, S., Kebukawa, Y., Kitadai, N., Igisu, M., & Matsuoka, N. (2018). Geochemistry and the Origin of Life: From Extraterrestrial Processes, Chemical Evolution on Earth, Fossilized Life’s Records, to Natures of the Extant Life. Life-Basel, 8(4). https://doi.org/ARTN3910.3390/life8040039

Payne, J. L., Boyer, A. G., Brown, J. H., Finnegan, S., Kowalewski, M., Krause, R. A., Lyons, S. K., McClain, C. R., McShea, D. W., Novack-Gottshall, P. M., Smith, F. A., Stempien, J. A., & Wang, S. C. (2009). Two-phase increase in the maximum size of life over 3.5 billion years reflects biological innovation and environmental opportunity. Proceedings of the National Academy of Sciences of the United States of America, 106(1), 24–27. https://doi.org/10.1073/pnas.0806314106

Perunov, N., Marsland, R. A., & England, J. L. (2016). Statistical Physics of Adaptation. Physical ReviewX, 6(2). https://doi.org/ARTN02103610.1103/PhysRevX.6.021036

Pinero, J., & Sole, R. (2018). Nonequilibrium Entropic Bounds for Darwinian Replicators. Entropy (Basel), 20(2). https://doi.org/10.3390/e20020098

Prigogine, I., & Nicolis, G. (1971). Biological order, structure and instabilities. Q Rev Biophys, 4(2), 107–148. https://doi.org/10.1017/s0033583500000615

Ragazzon, G., & Prins, L. J. (2018). Energy consumption in chemical fuel-driven self-assembly. Nat Nanotechnol, 13(10), 882–889. https://doi.org/10.1038/s41565-018-0250-8

Roden, E. E., & Jin, Q. (2011). Thermodynamics of microbial growth coupled to metabolism of glucose, ethanol, short-chain organic acids, and hydrogen. Appl Environ Microbiol, 77(5), 1907–1909. https://doi.org/10.1128/AEM.02425-10

Saadat, N. P., Nies, T., Rousset, Y., & Ebenhoh, O. (2020). Thermodynamic Limits and Optimality of Microbial Growth. Entropy (Basel), 22(3). https://doi.org/10.3390/e22030277

Schrödinger, E. (1967). What is life? the physical aspect of the living cell & Mind and matter. University P.

Schneider, E.D., Kay, J.J., 1994. Life as a manifestation of the second law of thermo-dynamics. Math. Comp. Model. 19, 25–48.

Sessions, A. L., Doughty, D. M., Welander, P. V., Summons, R. E., & Newman, D. K. (2009). The continuing puzzle of the great oxidation event. Curr Biol, 19(14), R567–574. https://doi.org/10.1016/j.cub.2009.05.054

Silow, E. A., & Mokry, A. V. (2010). Exergy as a Tool for Ecosystem Health Assessment. Entropy, 12(4), 902–925. https://doi.org/10.3390/e12040902

Soo, R. M., Hemp, J., Parks, D. H., Fischer, W. W., & Hugenholtz, P. (2017). On the origins of oxygenic photosynthesis and aerobic respiration in Cyanobacteria. Science, 355(6332), 1436–1440. https://doi.org/10.1126/science.aal3794

Spanner, D. C. (1953). Biological Systems and the Principle of Minimum Entropy Production. Nature, 172(4389), 1094–1095. https://doi.org/DOI10.1038/1721094a0

Ulanowicz, R. E. (2001). Information theory in ecology. Comput Chem, 25(4), 393–399. https://doi.org/10.1016/s0097-8485(01)00073-0

Walker, S. I. (2017). Origins of life: a problem for physics, a key issues review. Rep Prog Phys, 80(9), 092601. https://doi.org/10.1088/1361-6633/aa7804

Wu, Z. J., Wu, X. F., Yang, Z. H., & Ouyang, L. N. (2017). A simple thermodynamic model for evaluating the ecological restoration effect on a manganese tailing wasteland. Ecological Modelling, 346, 20–29. https://doi.org/10.1016/j.ecolmodel.2016.12.008

Wu, Z. J., Wu, X. F., Yang, Z. H., & Ouyang, L. N. (2018). Internal energy ratios as ecological indicators for description of the phytoremediation process on a manganese tailing site. Ecological Modelling, 374, 14–21. https://doi.org/10.1016/j.ecolmodel.2018.02.009

Yamamoto, M., & Takai, K. (2011). Sulfur metabolisms in epsilon-and gamma-proteobacteria in deep-sea hydrothermal fields. Front Microbiol, 2, 192. https://doi.org/10.3389/fmicb.2011.00192

Zeng, H., & Yang, A. (2020). Bridging substrate intake kinetics and bacterial growth phenotypes with flux balance analysis incorporating proteome allocation. Sci Rep, 10(1), 4283. https://doi.org/10.1038/s41598-020-61174-0

